# Walking kinematics in the polymorphic seed harvester ant *Messor barbarus:* influence of body size and load carriage

**DOI:** 10.1101/614362

**Authors:** Hugo Merienne, Gérard Latil, Pierre Moretto, Vincent Fourcassié

## Abstract

Ants are famous in the animal kingdom for their amazing load carriage performances. Yet, the mechanisms that allow these insects to maintain their stability when carrying heavy loads have been poorly investigated. Here we present a study of the kinematics of loaded locomotion in the polymorphic seed-harvesting ant *Messor barbarus*. In this species big ants have larger heads relative to their size than small ants. Hence, their center of mass is shifted forward, and the more so when they are carrying a load in their mandibles. We tested the hypothesis that this could lead to big ants being less statically stable than small ants, thus explaining their lower load carriage performances. When walking unloaded we found that big ants were indeed less statically stable than small ants but that they were nonetheless able to adjust their stepping pattern to partly compensate for this instability. When ants were walking loaded on the other hand, there was no evidence of different locomotor behaviors in individuals of different sizes. Loaded ants, whatever their size, move too slowly to maintain their balance through dynamic stability. Rather, they seem to do so by clinging to the ground with their hind legs during part of a stride. We show through a straightforward model that allometric relationships have a minor role in explaining the differences in load carriage performances between big ants and small ants and that a simple scale effect is sufficient to explain these differences.

## Introduction

The locomotion of animals can be described as a succession of strides that follows a specific inter-leg coordination pattern called gait (Alexander, 2003). In hexapod animals such as insects the most common gait is the alternating tripod (Delcomyn, 1981), in which the animal walks by alternating the movement of two distinct sets of legs (the ipsilateral front and hind leg and the contralateral mid leg, e.g. L1, L3, R2 and R1, R3, L2 respectively, with L for left and R for right), each of which forms a tripod supporting the body. In its ideal form, the two tripods perfectly alternate: all the legs in one tripod group simultaneously lift-off while all the legs of the other tripod group are still on the ground. However, depending on various features of their locomotion, insects can also adopt more complex gait. For example, the pattern of leg coordination can change with locomotory speed (Bender et al., 2011; Wosnitza et al., 2012; Mendes et al., 2013; Wahl et al., 2015), walking curvature (Zolliköfer, 1994a) and direction of movement, i.e. forward or backward movement (Pfeffer et al., 2016). Insects also adapt their gait according to the features of the terrain on which they are moving, e.g. when they walk on a non-level substrate (Seidl and Wehner, 2008; Reinhardt et al., 2009; Grabowska et al., 2012; Ramdya et al., 2017; Wöhrl et al., 2017) or when they climb over obstacles (Watson et al., 2002). Another factor that is known to have an effect on leg coordination during locomotion in terrestrial vertebrates (Jagnandan and Higham, 2018) but that has been less studied in insects is the change in the total mass an individual has to put in motion. Changes in total mass can be progressive or sudden and can occur in a variety of situations. For example, it happens in female insects during egg development and after oviposition, autotomy, i.e. the voluntary shedding of a body segment (Fleming and Bateman, 2007; Lagos, 2017), or, in the most common case, when insects are transporting food, either internally, after ingesting liquid, or externally, in their mandibles. In all these situations the change in total mass induces a shift in the center of mass of the insect which can profoundly affect its locomotion.

Ants offer a very good model to study the effect of changes in total mass on walking kinematics for at least three reasons. First, they are famous for their load carriage performances and can routinely carry loads (prey items, seeds, nest material, nestmates and brood) weighing more than ten times their own mass over tens of meters (Bernadou et al., 2016). In addition, the food they collect can be transported internally or externally. The shift in their center of mass can thus vary both in intensity and direction, which is likely to disrupt their walking kinematics in different ways. Second, due to the high number of species in their taxon (Hölldober & Wilson, 1990), the size and shape of ant bodies is extremely variable, which probably affects differently the kinematics of their locomotion. And third, ants live in very diverse environments and can be subterranean, ground-living, or arboreal (Hölldobler and Wilson, 1990), which is bound to constrain their movements and affect their locomotion differently (Gravish et al., 2013; Seidl and Wehner, 2008; Reinhardt et al., 2009; Wöhrl et al., 2017).

The effects of changes in total mass due to load carriage on the walking kinematics of ants have been poorly explored in the ant literature. The main effect of carrying a load in the mandibles is to induce a forward shift of the center of mass of the system formed by the ant and the load they carry. However, according to Hughes (1952), insects could counter this effect and achieve balanced locomotion by using static stability, i.e. by keeping the planar projection of their center of mass within the polygon formed by the legs simultaneously in contact with the ground (called polygon of support). In fact, this is what loaded ants do. For example, *Cataglyphis fortis* workers ensure static stability by placing their front legs in a more forward position when loaded than when unloaded and by reducing their stride length (Zollikofer, 1994b). In the species *Atta vollenweideri*, whose foraging workers carry elongated pieces of grass over their head, ants increase their mechanical stability by increasing the number of legs simultaneously in contact with the ground. They do so by increasing over consecutive steps the overlap between the stance (retraction) phase of the supporting tripod and some of the legs of the other tripod (mostly the front leg) and by dragging their hind legs during the swing (protraction) phase (Moll et al., 2013). These ants also adjust the angle of the load they carry so that the planar projection of their center of mass remains within the polygon of support (Moll et al., 2013).

In this paper, we studied the effect of load carriage on the walking kinematics of the seed-harvesting ant *Messor barbarus*, an ant species that is characterized by a highly polymorphic worker caste, i.e. a high variability in the size of individuals within the same colony. In addition, this polymorphism is characterized by allometric relationships between the different parts of the body (Bernadou et al., 2016), which means that big workers are not an enlarged copy of small workers but that the growth of some of their body parts during development is different from that of small workers (Bonner, 2006). In fact, relative to their mass their legs are shorter and their head bigger than those of small workers. Here, we hypothesized that this allometry could lead to differences in unloaded and/or loaded locomotion in different sized workers and thus could explain the differences observed in their load carriage performances (Bernadou et al., 2016). To test this, we ran an experiment in which we compared the kinematics of workers tested first unloaded and then loaded with loads whose relative mass we varied in a systematic way across different sized ants

## Materials and methods

### Studied species and rearing conditions

We used workers from a large colony of *M. barbarus* collected in April 2018 at St Hippolyte (Pyrénées Orientales) on the French Mediterranean coast. *Messor barbarus* is a seed harvester ant whose mature colonies can shelter several tens of thousands individuals (Cerdan, 1989). Its workers display a polymorphism characterized by a continuous monophasic allometry between head mass and thorax length (Heredia and Detrain, 2000; Bernadou et al., 2016). Individuals range from 1 to 40 mg in mass and from 2 to 15 mm in length. The colony was kept in a box (LxWxH: 0.50×0.30×0.15 m) with Fluon® coated walls to prevent ants from escaping. Ants nested inside test tubes covered with opaque paper. They had access *ad libitum* to water and to seeds of various species (canary grass, niger, oats). The experimental room was kept at a constant temperature of 28°C and under a 12:12 L:D regime.

### Experimental setup

The setup we used in our experiment was designed and built by a private company (R&D Vision, France. http://www.rd-vision.com). It consisted in a walkway (160 x 25mm) covered with a piece of black paper (Canson®, 160g/m^2^) on which the ants were walking during the experiment. The walkway was surrounded by five synchronized high speed video cameras (JAI GO-5000M-PMCL: frequency: 250Hz; resolution: 30µm/px for the top camera, 20μm/px for the others), one placed above the walkway and four placed on its sides (Figure 1). Four infrared spots (λ=850nm), synchronized with the cameras, were used to illuminate the walkway from above, allowing a better contrast between the ants and the background on the videos. The temperature on the walkway was monitored with an infrared thermometer (MS pro, Optris, USA, http://www.optris.com). Over the course of the experiment the mean temperature was (mean ± SD) 28 ± 1.4 °C.

**Figure 1:**
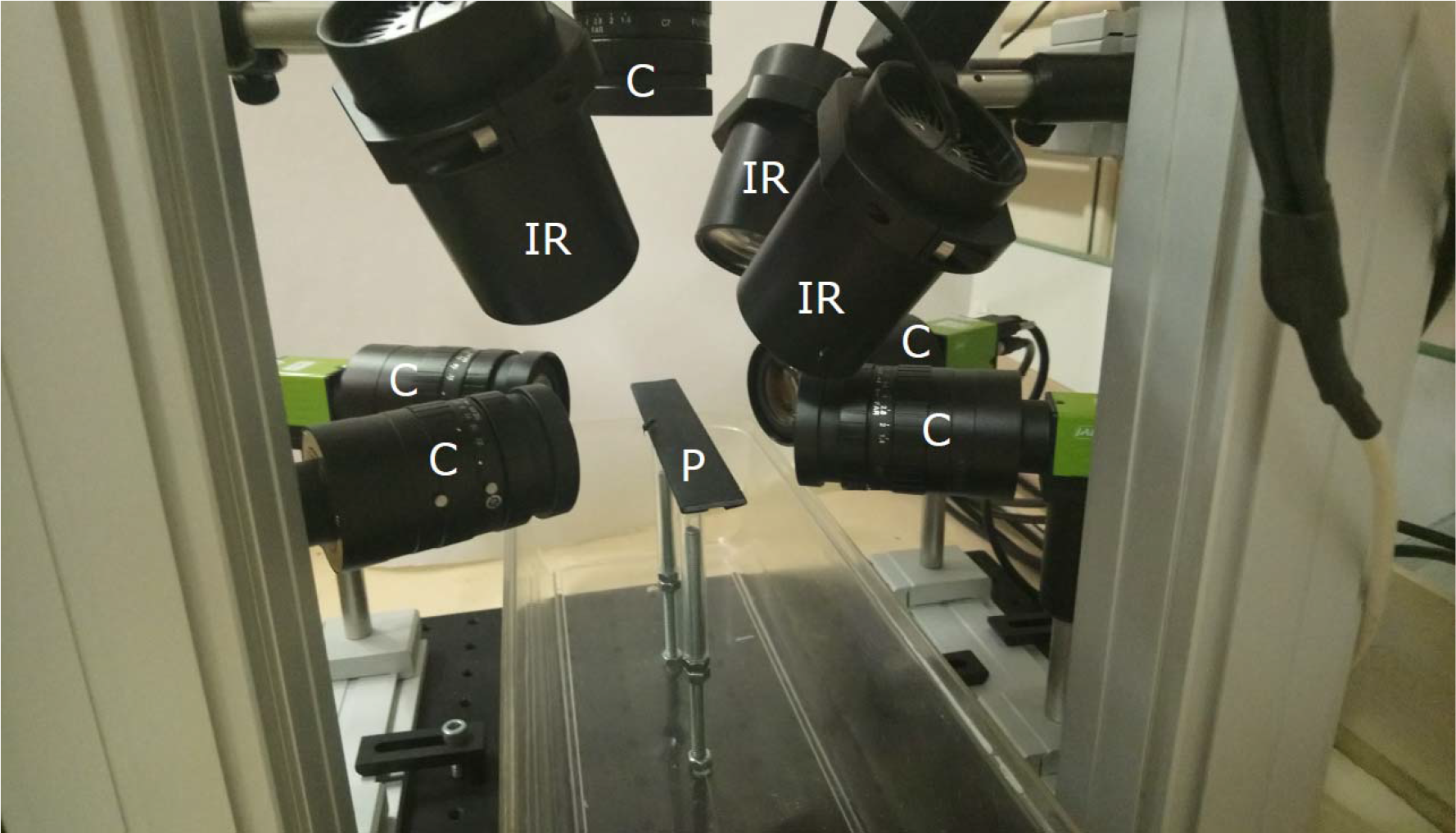
Video acquisition system. C: cameras; IR: infra-red spots; P: 25mm wide walkway.

### Experimental protocol

All experiments were performed between April and July 2018.

On the first day of a week in which we tested ants, we installed a bridge between the colony and a box in which a few seeds were placed. We then collected during one hour one ant out of three that carried a seed back to the colony. These ants were kept apart and used for the experiment in the following days. We also collected a few ants (weighing between 10-15 mg) to dissect their Dufour gland in order to create an artificial pheromone trail in the middle of the walkway (Heredia and Detrain, 2000). Since ants tended to follow the trail this increased the chance to obtain a straighter path in the middle of the walkway, which allowed us to neglect the effect of path curvature on ant kinematics (Zolliköfer, 1994a). In order to extract the Dufour gland, ants were first anesthetized by placing them in a vial plunged in crushed ice, then killed by removing their head and fixed on their back with insect pins under a binocular microscope. Their gaster was opened transversally with a scalpel following the first sternite and the ventral part was pulled away. The poison gland and the fat gland were then gently removed until the Dufour gland became visible. This latter was then collected and placed in a hexane solution to extract the trail pheromone. We used a concentration of 1 gland / 20μl which has been shown to be sufficient to elicit a clear trail following response in *M. barbarus* workers (Heredia and Detrain, 2000).

Each time an ant was tested, it was picked from the group of ants that had been separated on the first day of the week, then weighed and isolated in a small box with access to water. The ant was first tested unloaded and then loaded with lead fishing weights whose mass ranged from 2 to 100mg. Every five tested ant we made an artificial trail on the walkway by depositing every centimeter with a 10 μl syringe a droplet of 1 μl of the solution of Dufour gland extract. To reduce stress, ants were transferred from their individual box to the walkway by letting them climb up and down on a pencil. Once on the walkway, the movement of the ant was recorded as soon as it started to walk along a more or less straight path. The ant was then captured at the end of the walkway and anesthetized by placing it in a vial plunged in crushed ice. It was then fixed dorsally with Plasticine under a binocular microscope with its head maintained horizontally. This allowed us to put a drop of superglue (Loctite, http://www.loctite.fr) on the top of its mandibles and to glue a fishing weight. The same procedure as for unloaded ants was then used to film loaded ants. At the end of the experiment the ant was killed and we weighed its head, thorax (with the legs), and gaster separately to the nearest 0.1 mg with a precision balance (NewClassic MS semi-micro, Mettler Toledo, United Sates). These measures were used to compute the displacement of the center of mass (CoM) of the ants on the videos (see SI for details of the procedure).

Whether unloaded or loaded, we filmed all ants for at least three strides. We defined a stride period as the time elapsed between two consecutive lift off of the right mid leg. For our analysis, we cropped our videos to a whole number of strides.

### Data extraction and analysis

In order to compute the horizontal movement of the ants’ main body parts (gaster, thorax, head or head and load if one was carried, see SI for details) and of its overall center of mass (CoM) we used the software Kinovea (version 0.8.15, https://www.kinovea.org) to semi-automatically track points of interest on the videos of the top camera (Figure 2). When several video shootings of the same ant had been made, we selected the video in which the ant had the straighter path. As a criteria for path straightness we calculated the ratio of the distance actually traveled by the ant (based on the horizontal trajectory of its center of mass) on the straight line distance between the first and last point of the trajectory and considered that the path was sufficiently straight when this ratio was lower than 1.2.

**Figure 2:**
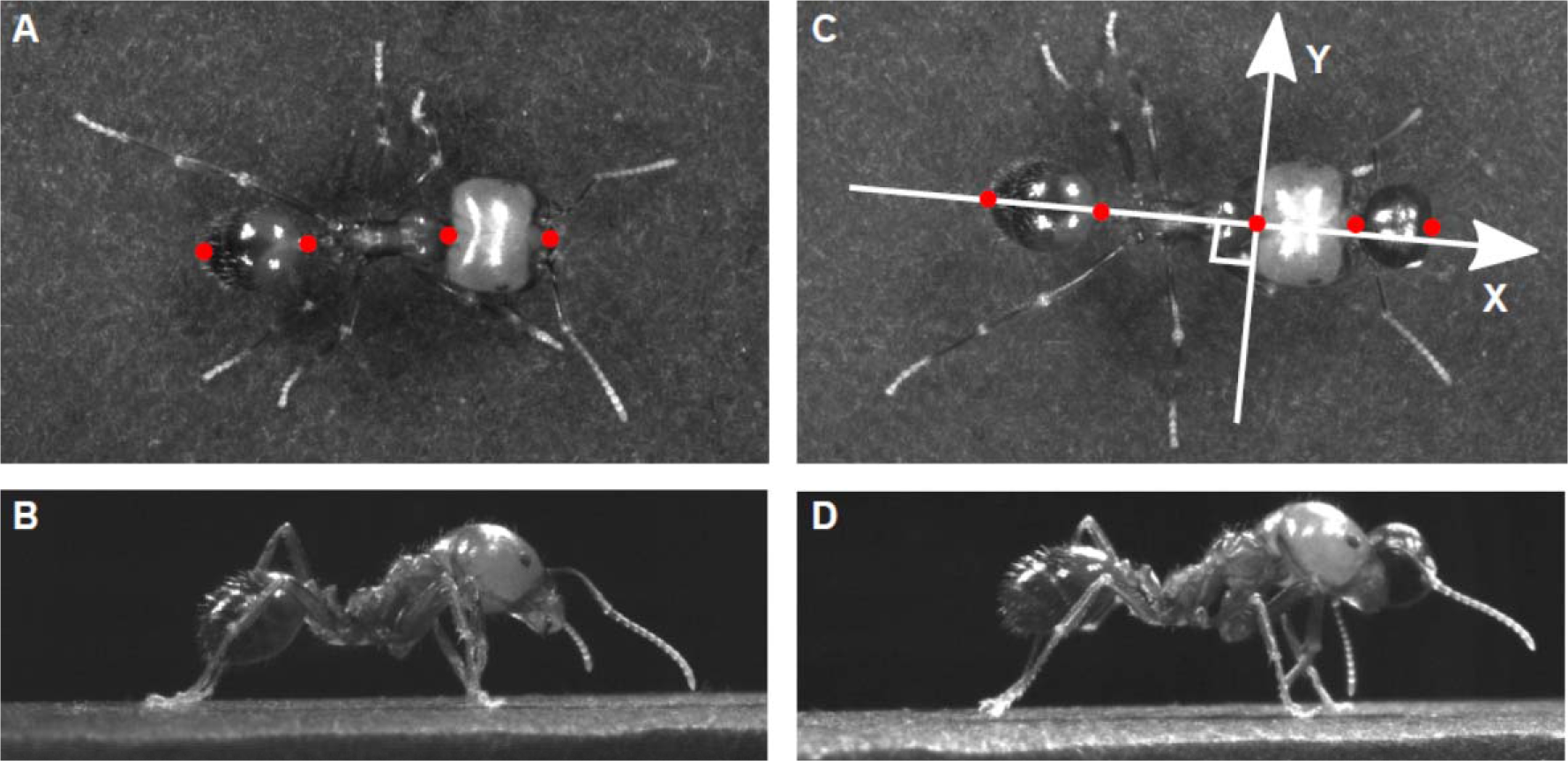
Location of the points tracked on each ant. The snapshots show a top view (**A**, **C**) and a side view (**B**, **D**) of the same ant (ant mass = 32.5mg) tested in unloaded (**A**, **B**) and loaded condition (**C**, **D**) (load mass = 63mg). In **C**) the X axis corresponds to the longitudinal body axis while the Y axis corresponds to the transverse body axis. The position of the track points are represented in red (See SI for details).

Using the software Kinovea and ImageJ (version 2.0.0, FIJI distribution, https://fiji.sc/) we then accessed the information corresponding to the stepping pattern of the ant. For each video frame we visually determined the state of each leg during locomotion (i.e. in stance phase, swung or dragged) on the lateral view of the ant and recorded the spatial position of the leg tarsus during the stance phases on the dorsal view. These positions were expressed in a coordinate system centered on the neck of the ant, with the X axis corresponding to the longitudinal axis of its body and the Y axis to the transverse axis (Figure 2). In order to compare ants of different sizes, all distances were normalized to the body length of the ant (calculated from the tip of the gaster to the tip of the mandibles).

We computed the duty factor for each leg as the fraction of the stride the leg was in contact with the ground (Ting et al., 1994; Spence et al., 2010). For each leg, we also computed the mean relative position at lift off (Posterior Extreme Position: PEP) and at touch down (Anterior Extreme Position: AEP) by averaging the relative positions of the leg over the strides we filmed. Since the path followed by the ant was straight, we also averaged the values of the right and left leg of each pair of legs when computing the duty factors and relative leg positions. Following Wosnitza et al. (2012) and Wahl et al. (2015) we calculated step amplitude rather than stride length. For each leg we computed step amplitude by averaging the distances between PEP and AEP positions in the ant coordinate system. Again, because the path followed by the ant was straight, we averaged the values of the right and left leg of each pair of legs when computing step amplitude.

We studied inter-leg coordination by comparing the time of lift off of every leg to the time of lift off of the right mid leg (Wosnitza et al., 2013; Wahl et al., 2015). More precisely, we computed, for each leg, the time lag between the leg lift off and the last lift off of the right mid leg. We then divided the value of the time lag by the time elapsed between two successive lift off of the right mid leg. This value was expressed as a phase shift between – π and π for each leg. Finally, we used circular statistics (Batschelet, 1981) to compute the mean vector of the distribution of the phase shifts for specific groups of ants. As an indication of how ant gait was close to an ideal alternating tripod locomotion we also computed the Tripod Coordination Strength (TCS) (Wosnitza et al., 2012; Wahl et al., 2015; Ramdya et al., 2017). This index can take values between 0 and 1. A TCS of 1 corresponds to a perfect alternating tripod gait while a TCS of 0 means that the ant performed a completely different type of gait.

Finally, following Moll et al. (2013), we also computed for each ant the change over time of the static stability margin (SSM) during locomotion. For every video frame, the SSM was defined as the minimum distance between the horizontal projection of the ant CoM and the edges of the polygon formed by all legs in contact with the ground, including the dragged legs. The SSM is positive if the projection of the CoM lies inside the polygon, negative otherwise. We considered that the locomotion was statically stable when the ant managed to maintain static stability during locomotion (i.e., when the SSM was positive) and that it was statically unstable when it was not the case (i.e., when the SSM was negative).

We performed all data analysis and designed all graphics with R (version 3.5.1) run under RStudio (version 1.0.136). For unloaded condition, we expressed all kinematic variables as a linear function of the decimal logarithm of ant mass. For loaded condition, because the same ants were tested loaded and unloaded, we calculated the difference in the value of each kinematic variable between loaded and unloaded conditions and expressed it as a linear function of both the decimal logarithm of ant mass and the decimal logarithm of load ratio which was defined as 1 + (load mass/ant body mass) (Bartholomew et al., 1988).

## Results

### Unloaded ants: influence of body mass (Table 1)

Stride frequency (*F*_1,43_ = 64.82, *P*<0.001) and step amplitude for every leg (front leg: *F*_1,43_ = 4.1, *P*=0.049; mid leg: *F*_1,43_ = 13.0, *P*<0.001; hind leg: *F*_1,43_ = 30.0, *P*<0.001) decreased with ant mass. As a result, the speed decreased with ant mass as well (*F*_1,43_ = 109.2, *P*<0.001; Figure 3). Thus, relative to their size, big ants move more slowly than small ants. However, absolute speed did not depend on ant mass (mean ± SD: 29.1 ± 4.5 mm.s^−1^)

**Figure 3:**
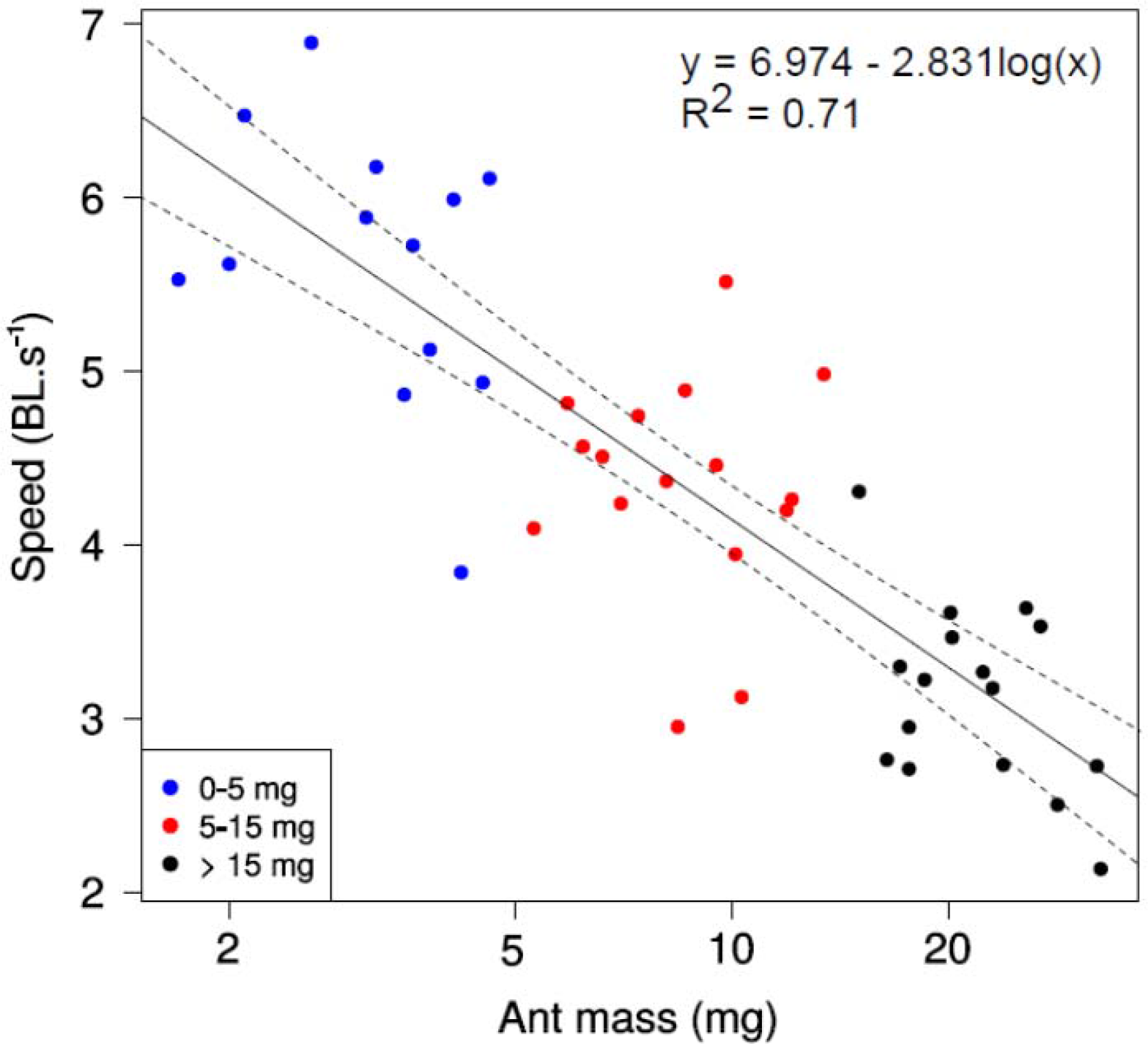
Speed as a function of ant mass for unloaded ants. The straight line gives the prediction of a linear regression model (F1,43 =115.7, P<0.001) and the dashed lines gives the 95% confidence interval of the slope of the regression line (N= 45 ants).

The duty factor of all legs increased with increasing ant mass, particularly for the front (*F*_1,43_ = 36.7, *P*<0.001) and mid (*F*_1,43_ = 22.6, *P*<0.001) legs (Figure S1A-C). Therefore, compared to small ants big ants tend to have more legs in contact with the ground during a stride (*F*_1,43_ = 36.4, *P*<0.001). The front and mid legs were almost never dragged by the ants. However, independent of ant mass, hind legs were dragged during 23 % of a stride on average.

In an ideal alternating tripod gait, all legs of a tripod lift off simultaneously. In actual locomotion however, there is no such perfect synchronization. Nevertheless, the alternating tripod gait model still holds if the time interval between the lift off of the three legs of the same tripod is small. Figure 4A shows that the ants’ gait is very close to an ideal tripod gait (see also Fig. S2A). However, for bigger ants, the front legs tended to lift off slightly after the mid and hind legs of the same tripod. As a result, the TCS slightly decreased for bigger ants (*F*_1,43_ = 6.3, *P*= 0.016).

**Figure 4:**
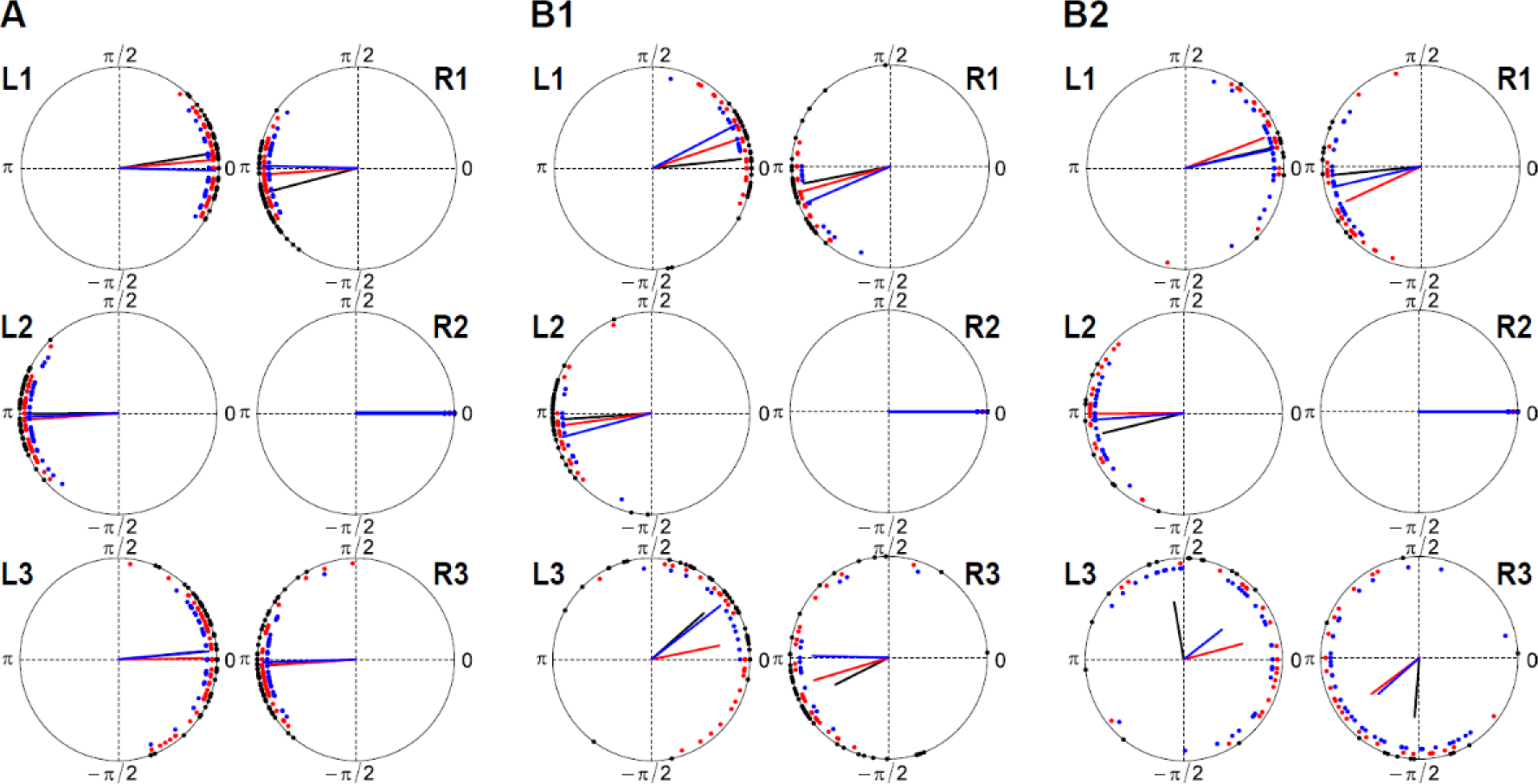
Phase plots of lift off onset of all legs with respect to the right mid leg (R2) **A** unladen ants (N_0-5mg_ = 13; N_5-15mg_ = 16; N_>15mg_ = 15); **B** – laden ants; **B1** – Load Ratio ≤ 3.5 (N_0-5mg_ = 4; N_5-15mg_ = 9; N_>15mg_ = 12)); **B2** – Load Ratio > 3.5 (N_0-5mg_ = 9; N_5-15mg_ = 7; N_>15mg_ = 3). R, right; L, left; blue: data for small ants (0-5mg); red: data for intermediate ants (5-15mg); black: data for big ants (>15mg); lines: mean vector – length is inversely proportional to dispersion.

The front legs tended to be positioned at a larger distance from the longitudinal body axis (Y position) in big ants compared to small ants both at touch down (AEP) (*F*_1,43_ = 17.5, *P*<0.001) and lift off (PEP) (*F*_1,43_ = 15.5, *P*<0.001). All legs, especially the hind legs, tended to be positioned in a more forward position (X position) at lift off (PEP) in big ants compared to small ants (front legs: *F*_1,43_ = 20.4, *P*<0.001; mid legs: *F*_1,43_ = 16.6, *P*<0.001; hind legs: *F*_1,43_ = 50.5, *P*<0.001) (Figure 5A).

**Figure 5:**
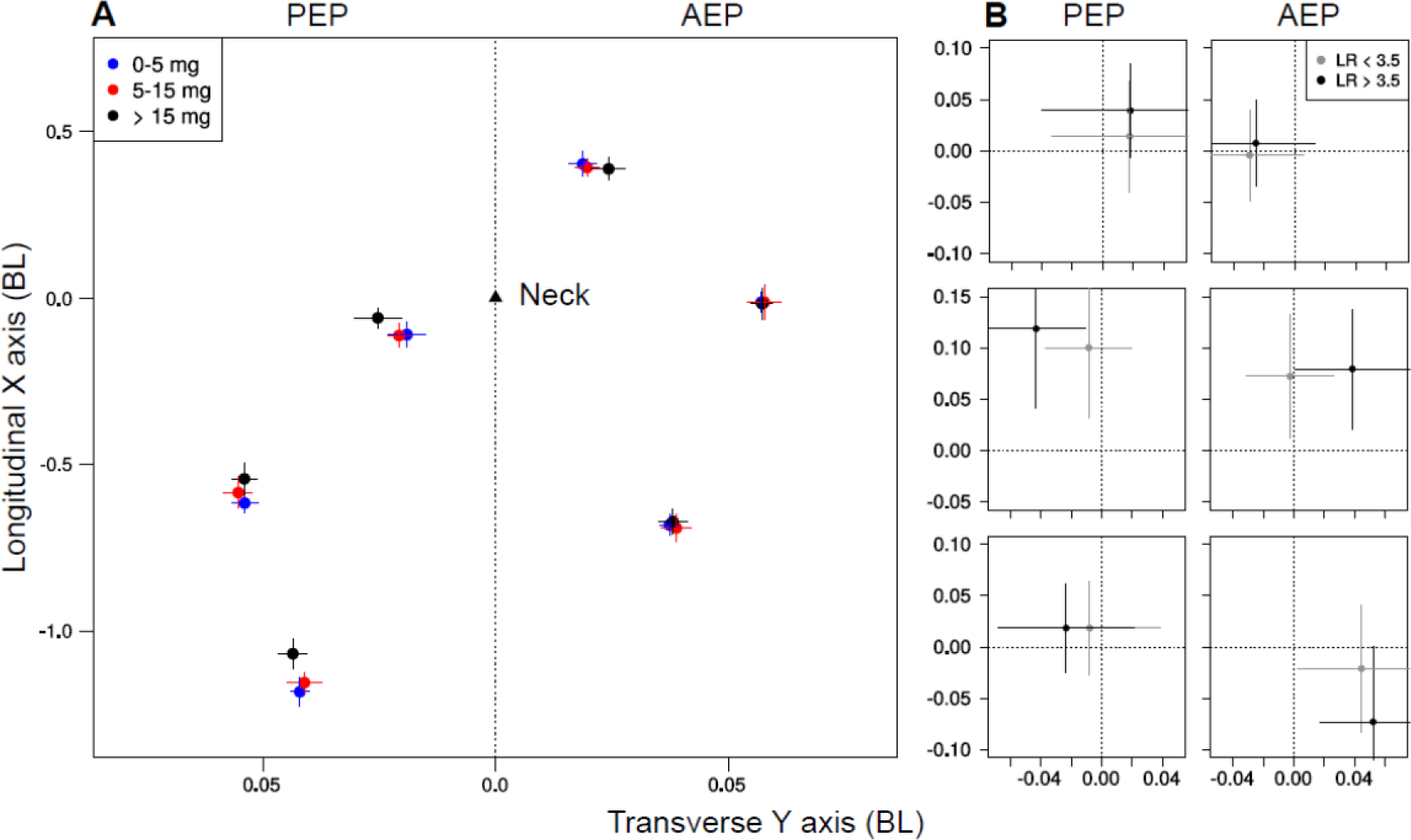
Footfall geometry of ants during locomotion. **A**: Unloaded ants; the mean position of the front, mid, and hind legs during lift off (PEP) and touch down (AEP) along with their standard deviation is shown for different ranges of ant body masses (N_0-5mg_ = 13, N_5-15mg_ = 16, N_>15mg_ = 16). **B**: Loaded ants; changes in leg positions at lift off (PEP) and touch down (AEP) when ants were walking loaded compared to when they were walking unloaded. The origin corresponds to the leg position for unloaded ants. The average change in position over three strides along with their standard deviation is shown. Ants were categorized in two groups depending on load ratio (N_LR<3.5_ = 26, N_LR>3.5_ = 19). The scale is in units of body length.

The static stability margin decreased during a stride and reached a local minimum value just before touch down of one of the front legs (Figure 6A & 6B). The minimum value of the static stability margin decreased with increasing ant mass (*F*_1,43_ = 4.7, *P*= 0.036). Moreover, the proportion of time an ant moved in statically unstable locomotion increased with ant mass (*F*_1,43_= 5.0, *P*= 0.030, compare Figures 6A & 6B). Therefore, small ants have a more balanced locomotion than big ants.

**Figure 6:**
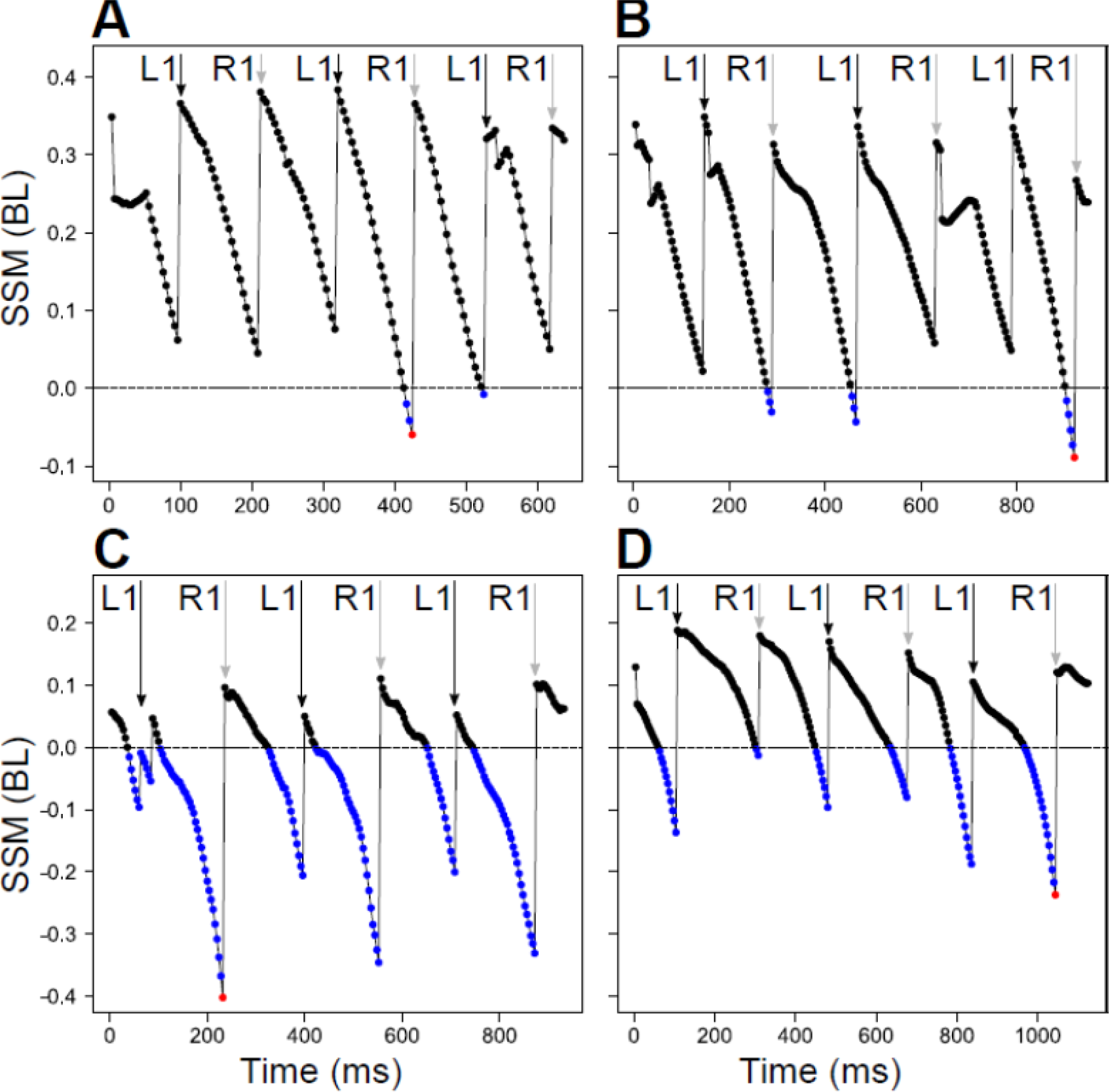
Time variation of the Static Stability Margin (SSM). The value of SSM for unloaded (**A-B**) and loaded (**C-D**) ants, normalized by body length, is shown during three consecutive strides for a small ant (**A-C**) (ant mass = 4.2 mg, LR = 5.0) and a big ant (**B-D**) (ant mass = 32.1 mg, Load Ratio = 2.0). Black and grey arrows represent R1 and L1 touched down, respectively, blue dots correspond to negative SSM and red dots correspond to the minimum SSM.

### Loaded ants: influence of ant body mass and load ratio (Table 2)

Figure 7 shows the values of load ratio tested for ants of different body masses. Depending on the value of the load ratio, ants exhibited two kinds of behaviors when loaded. They could either keep the load lifted above the ground during locomotion or they could maintain their head in a very slanted position and push the load in front of them (see SI Movies 1-3 for illustrations). We called the first behavior “carrying” and the second “pushing”. The criteria we used to distinguish between the two behaviors is based on whether or not the load glued on the ant mandibles was in contact with the ground during locomotion. Pushing generally occurred for load ratio higher than five for ants above 10mg, while for ants of lower body mass both carrying and pushing could be observed for load ratio higher than four (Figure 7). We will only consider ants that carry their load in the following analyses.

**Figure 7:**
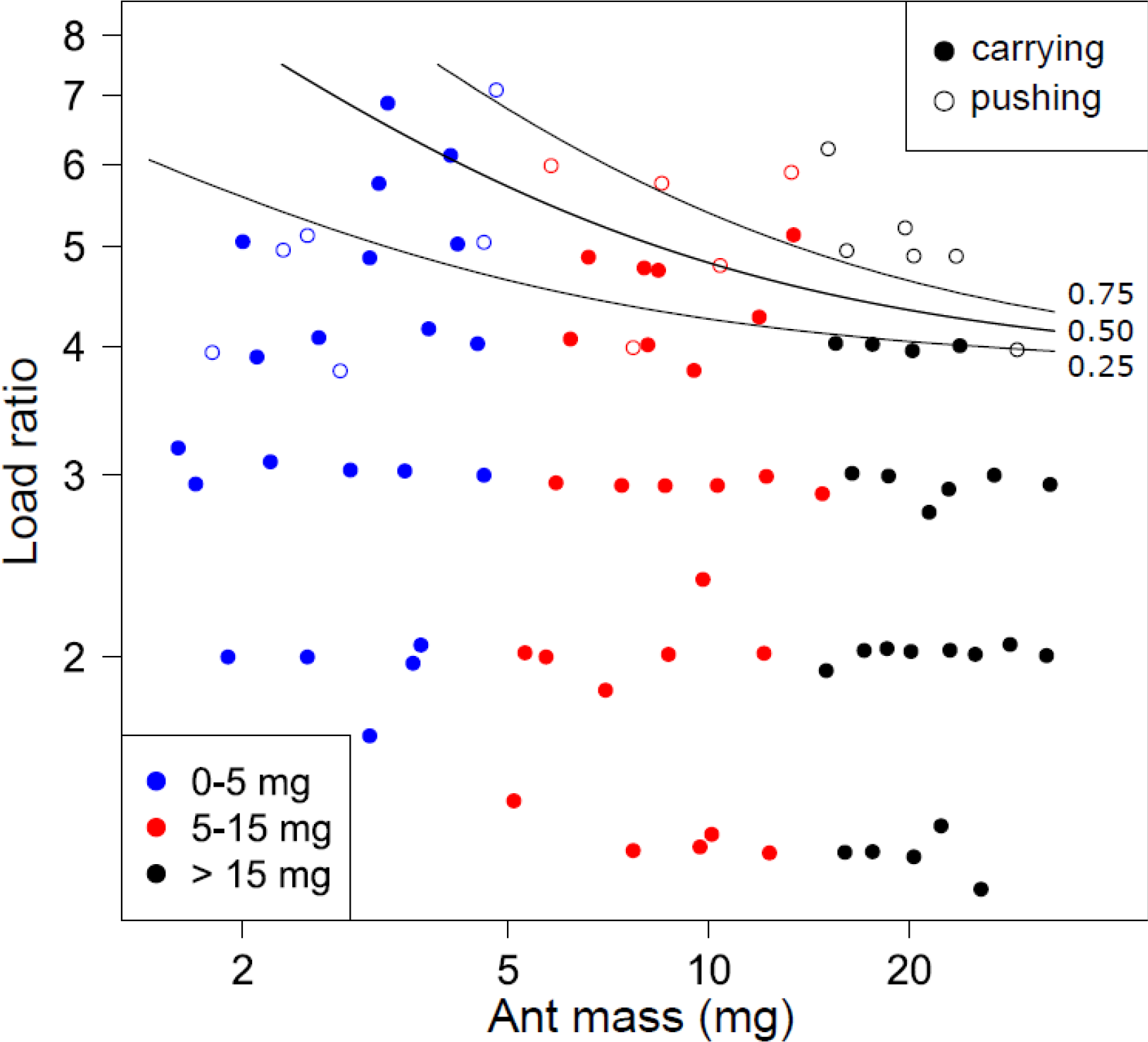
Transportation method used by ants during locomotion. Probability of pushing a load as a function of ant mass and load ratio. The lines of equal probability were calculated by a logistic regression model (N = 86 ants).

Independent of ant mass, stride frequency decreased with increasing load ratio (*F*_2,42_ = 19.7, *P*<0.001). Step amplitude was independent of load ratio for front and mid legs but decreased for hind legs (*F*_2,42_ = 3.2, *P*=0.051). Consequently, ant speed decreased with increasing load ratio (*F*_2,42_ = 23.6, *P*<0.001). However, for ants transporting equivalent loads there was no effect of body mass on both stride frequency and step amplitude.

Independent of ant mass, the duty factor increased for the front (*F*_2,42_ = 62.2, *P*<0.001), mid (*F*_2,42_ = 27.1, *P*<0.001) and hind legs (*F*_2,42_ = 14.9, *P*<0.001) for increasing load ratio and, independent of load ratio, the duty factor increased with ant mass, confirming the results obtained on unloaded ants (Figure S1D-F). The mean number of legs simultaneously in contact with the ground increased for increasing load ratio independent of ant size and, to a lesser extent, increased for increasing body mass independent of load ratio (*F*_2,42_ = 57.7, *P*<0.001). Similar to what occurred when ants were unloaded, the front and mid legs were almost never dragged during locomotion when ants were loaded.

When the ants were walking loaded their mid and hind legs tended to be more distant from their longitudinal body axis (Y position) with increasing load ratio, both during lift off (mid legs: *F*_2,42_ = 10.0, *P*<0.001; hind legs: *F*_2,42_ = 5.3, *P*= 0.009) and touch down (mid legs: *F*_2,42_ = 6.1, *P*=0.005; hind legs: *F*_2,42_ = 5.5, *P*= 0.007) (Figure 5B).

While performing loaded locomotion, ants did not exhibit the same inter-leg coordination pattern than during unloaded locomotion (Figure 4). First, there was more dispersion in phase shift between legs for loaded ants, especially for the hind legs and for high values of load ratio (> 3.5, see Figure 4B2). Second, the three legs of the same tripod tended to lift off in a specific order (i.e. mid leg -> front leg -> hind leg). This was especially clear for the hind leg, which was the last to lift off in each tripod. This order seems to be more strictly followed for higher load ratio and for bigger ants (Figure 5B2). As a result, the value of TCS decreased for increasing load ratio (*F*_2,42_ = 8.7, *P*<0.001).

Independent of ant size, the proportion of time ants were performing statically unstable locomotion increased with increasing load ratio in loaded ants (*F*_2,42_ = 65.4, *P*<0.001).

## Discussion

In this study, we investigated the kinematics of locomotion of unloaded and loaded ants of the polymorphic species *M. barbarus*. We found that, relative to their size, small ants were able to carry larger loads than big ants. Small ants also walked faster and were more stable than big ants; all ants, whatever their size, reduced their speed when carrying loads of increasing mass. The locomotion of unloaded ants was very close to an ideal alternating tripod gait. This allowed them to perform a rather statically stable locomotion. On the other hand loaded ants were most of the times statically unstable and their gait changed to more tetrapod-like locomotion, wave gait locomotion and hexapodal stance phases (Figure S2B).

### Unloaded ants

In *M. barbarus* big ants have, relative to their size, bigger heads than small ants (Heredia and Detrain, 2000; Bernadou et al., 2016). This means that their center of mass is located in a more anterior position compared to small ants. Big ants are thus more likely to be off balance than small ants. Therefore, one should expect static stability to decrease with increasing ant mass, which is what we actually observed. Nevertheless, the question arises of whether the decrease in stability we observed is the same as the decrease one should observed mechanically because of the forward shift of the center of mass of the body, or whether this decrease is less than the one expected because big ants adjust their gait in order to maintain their stability. In order to answer this question we modeled an ideal tripod gait for ants of different sizes. Our model took into account the difference in morphology across ants (for both mass repartition between body parts and relative leg lengths) but assumed ants had the same stepping pattern (based on the mean value of the leg positions observed in our experiment for all ants and then corrected for leg length, see SI for details). Following the method described in the method section, we then computed the minimum static stability margin (SSM) and the proportion of statically unstable locomotion for these ants. The slope of the regression line describing the relationship between the minimum static stability margin and log_10_(ant mass) and the proportion of statically unstable locomotion and log_10_(ant mass) was −0.142 and 0.183, respectively. These values should be compared to the values we found in our experiment, i.e., (mean ± CI_0.95_) −0.041 ± 0.036 and 0.029 ± 0.026, respectively (see Table 1). Thus, if ants were walking with an ideal tripod gait, the influence of ant mass would be more important than what we observed in our data. This means that big ants adjust their locomotion in order to increase their static stability. That this is indeed the case is suggested by the differences observed between small and big ants in both leg positioning and in gait parameters (Table 1). In big ants the front legs lifted off in a more anterior position, so that the minimum SSM (which occurs just before the front leg touch down, Figure 6) was less critical. This led to both an increase in duty factor and a decrease in step amplitude for front and mid legs. In conclusion, the differences in morphology between ants of different sizes do induce a less statically stable locomotion in big ants but this effect is reduced by the fact that they are able adjust their stepping pattern to compensate for this instability.

**Table 1:**
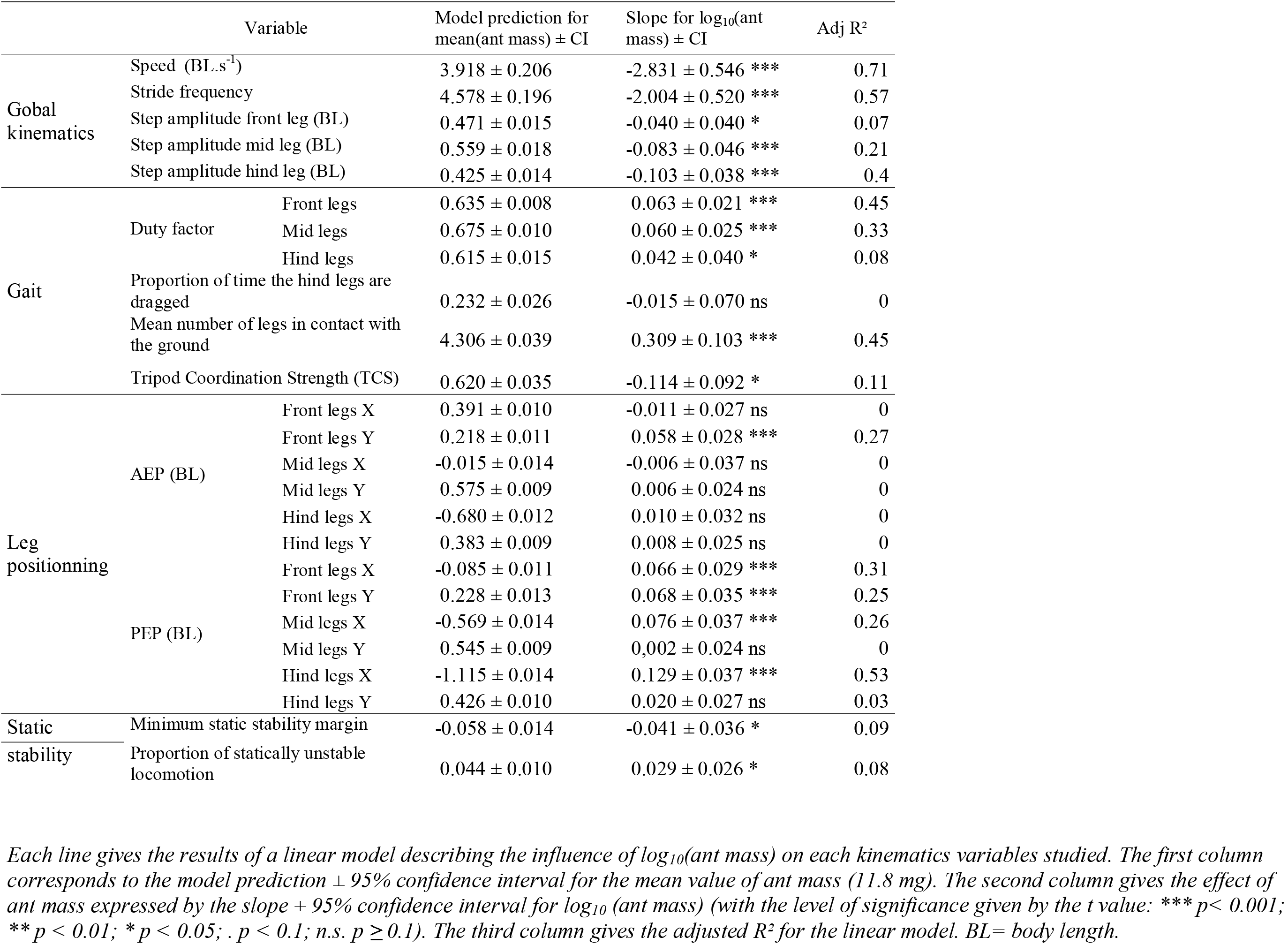
Influence of body mass (in mg) on the kinematics of unloaded ants (N = 45).

### Carrying capacity

Small ants were able to carry relatively heavier loads than big ants. This is concordant with the results obtained by Bernadou et al. (2016) in the same species and with those obtained by Burd (2000) in the leaf cutting ants *Atta colombica and A. cephalotes*. This difference in load carrying capacity can be accounted for by two non-exclusive explanations. The first is related to the well-known scale effect, while the second is related to differences in the locomotion and/or the morphology (induced by allometric relationships) of ants of different sizes.

The scale effect is due to the fact that the section of the muscles of an animal (which is directly related to the mechanical power they can develop) increases as the square of its body length while body mass increases as the cube (Schmidt-Nielsen, 1984; Dial et al., 2008). This would lead to a reduction in relative load capacity in big ants compared to small ants. However, this reasoning would hold only if big ants were a simple enlargement of small ones, i.e. if their body parts grew isometrically. As mentioned before, this is not exactly the case in *M. barbarus*: compared to small ants, big ants not only have larger heads (Bernadou et al., 2016, Fig. S5) but they have also relatively shorter legs (Fig. S6). Nonetheless, the scale effect could still apply to some extent. To assess its importance we compared our data of load carrying capacity in ants of different sizes to those that would be expected if the predictions of the scale effect were computed on ants of different sizes but with same morphology. We considered as the basis for the computation of our predictions the carrying capacity we observed for an ant weighing 1.5mg. As can be seen in Figure 8, the predictions of the carrying capacity for ants of different sizes is close to the curve based on our experimental data representing the 50% probability of carrying a load vs pushing it. It is also close to the curve representing the 50% probability of carrying a load vs dragging it from the field experiments by Bernadou et al. (2016) in which the ants transported food items of various sizes deposited on their foraging trails. Therefore, it seems that ants start pushing in our experiment for about the same load ratio values as they start dragging in Bernadou et al. (2016). Pushing instead of dragging probably occurs as a result of the load being glued on the mandibles.

**Figure 8:**
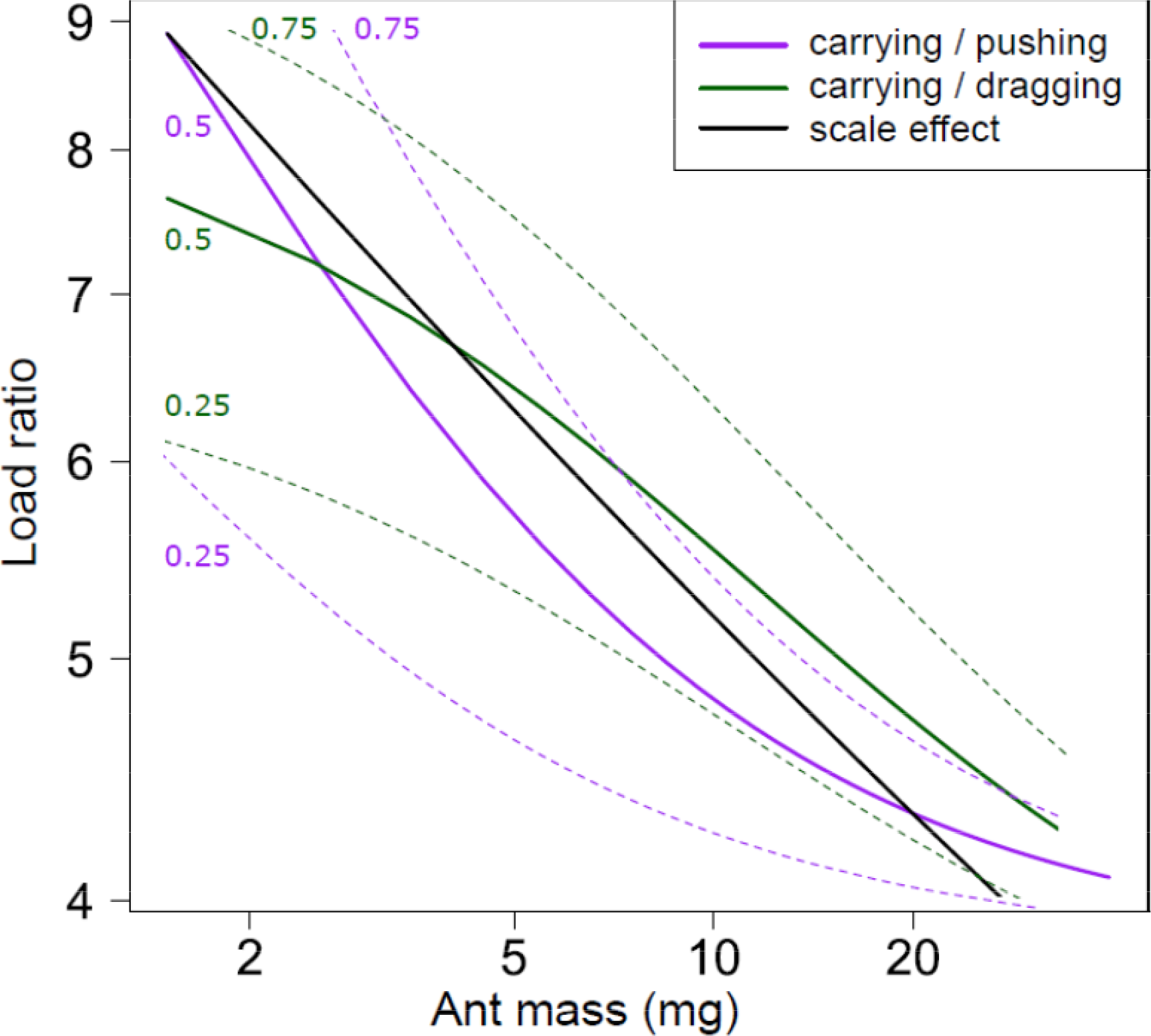
Ant carrying capacities and scale effect prediction. The purple line represents 50% probability of carrying the load versus pushing it; The green line represents 50% probability of carrying the load versus dragging it (data from (Bernadou *et al.*, 2016)); in both cases, the dashed lines represent 25% and 75% probabilities; the black line represents the prediction of load carrying capacity based on scale effect and a 1,5mg ant reference.

Nonetheless, one cannot exclude that the differences observed in locomotor behavior between small and big ants could also partly explain the differences in carrying capacity. Table 2 indeed points out some differences in the kinematics of ants of different sizes, independent of load ratio. However, these differences do not follow any particular logic and are difficult to interpret. Most of the observed differences in carrying capacities in ants of different sizes can thus be explained by a scale effect.

**Table 2:**
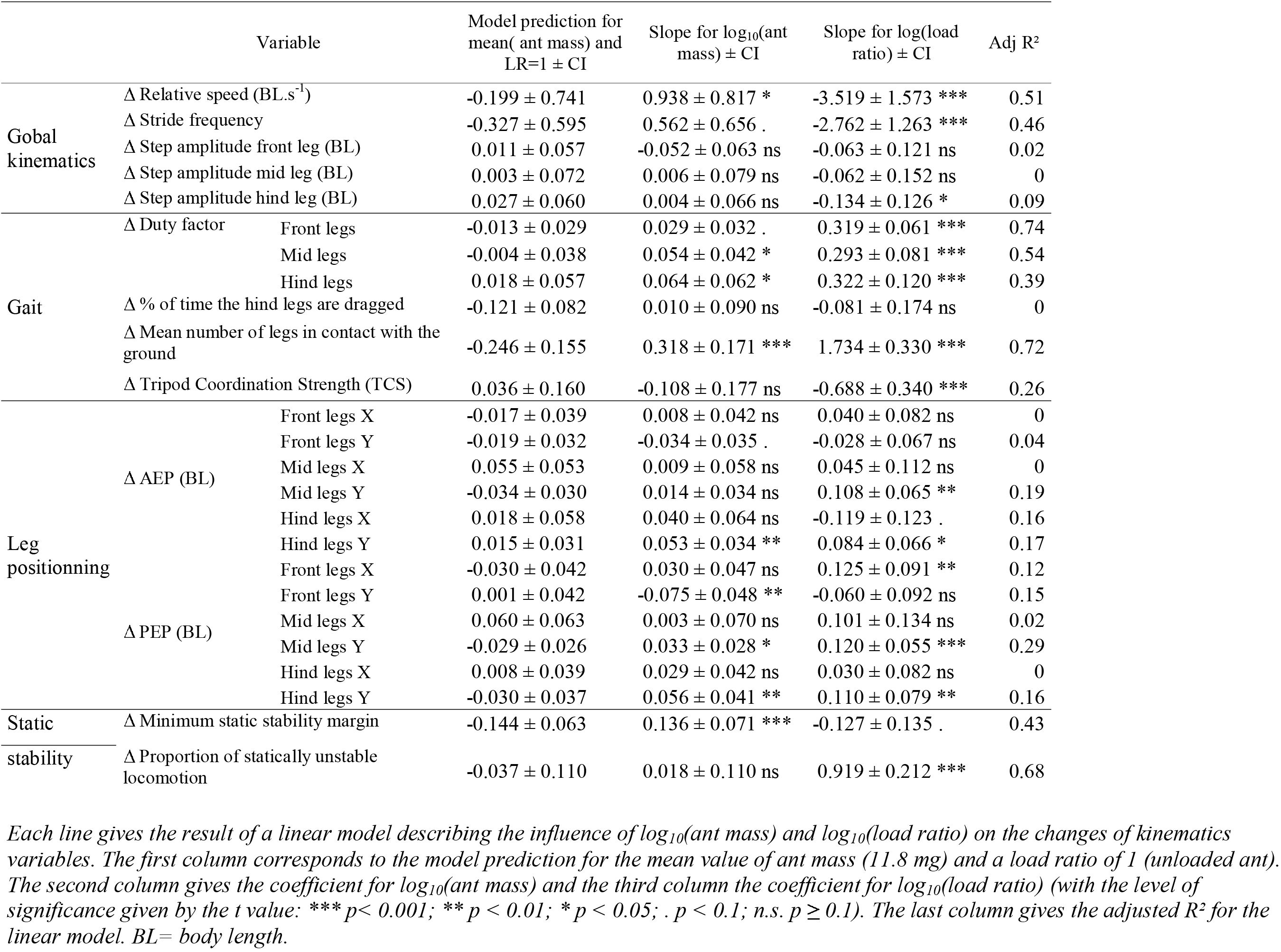
Influence of body mass (mg) and load ratio on the changes in kinematics between unloaded and loaded locomotion (N = 45).

### Influence of load ratio on locomotion

The main effect of carrying a load for an ant is to shift its CoM forward. As a consequence the CoM is located closer to the front edge of the polygon of support, or even lies out of it, and the SSM decreases or becomes negative, leading the ant to perform less statically stable or statically unstable locomotion. Moll et al. (2013) showed that loaded *Atta vollenweideri* ants can reduce this effect by changing the way they carry their load: by carrying the pieces of grass blade they hold in their mandibles in a more upward, backward-tilted position they can shift their CoM in a somewhat backward position. This is of course impossible in our experiment because ants cannot adjust the position of the load glued on their mandibles. This would not happen either in the field because most of the seeds collected by *M. barbarus* are not elongated enough to be carried in the same way as pieces of grass blades in grass-cutting ants. The CoM of loaded *M. barbarus* workers is thus shifted forward and the proportion of time their locomotion is statically unstable during a stride increases for increasing load ratio (Table 2), reaching up to 90% for the highest load ratios.

Such statically unstable locomotion has already been reported for insects in the literature. For instance, the cockroach *Blaberus discoidalis*, when moving at very high speed, often performs statically unstable locomotion and thus maintains its balance through dynamic stability (Ting et al., 1994; Koditschek et al., 2004). Dynamic stability refers to individuals keeping their balance when statically unstable by only briefly “falling” forward before new supporting legs contact the ground (Moll et al., 2013). Statically unstable locomotion has also been observed by Moll et al. (2013) in loaded workers of the grass cutting ant *A. vollenweideri*. These authors have suggested that loaded ants could use dynamic stability in order to avoid falling over during the statically unstable part of their locomotion (Moll et al., 2010, 2013).

However, loaded *M. barbarus* workers move too slowly (Table 2) to maintain their balance through dynamic stability: they would fall forward before the front leg catches up. Rather, we assume that they maintain their balance by clinging to the ground with the tarsal claws located at the end of their mid and hind legs. Consequently, they tend to keep more legs in contact with the ground for increasing load ratio. This leads to a decrease of their stride frequency and TCS and to an increase of the duty factor of all legs (Table 2). Hind legs are of particular importance in keeping the ant balanced because they have a higher lever-arm effect. In our experiment the percentage of time the hind legs were dragged decreases as soon as the ant was loaded independent of ant mass and load ratio (mean ± CI_0.95_: −12,1 ± 8,2 %), probably because, due to the position of the claws on the pretarsus (Fig; S3), the hind legs can better cling to the ground when they are not moving. In this respect, it would be interesting to investigate how ants maintain their balance when their adhesive prestarsal structures are blocked or when they walk on a slipping substrate (see Ramdya et al., 2017 for an example in *Drosophila melanogaster*). The tendency for the hind legs to lift off after the front legs touched down also increased for increasing load ratio (Figure 4B1 & 4B2). This is coherent with the balance strategy used by ants, as the SSM is maximal at front leg touch down (Figure 6C & 6D) and thus it is less risky to lift off the hind leg at this time. Finally, as a result of the stride frequency diminution (and because step amplitude remains constant), the speed decreases with increasing load ratio, which is concordant with most studies in other load carrying ants, e.g. *Atta colombica* (Lighton et al., 1987), *A. vollenweideri* (Röschard and Roces, 2002) and *Veromessor pergandei* (Rissing, 1982).

Reinhardt and Blickhan (2014a) showed that, during steady state locomotion, *Formica polyctena* uses mainly its hind legs in order to generate propulsion forces while Wöhrl et al., (2017) showed in *Cataglyphis fortis* that it is the mid legs that are mainly used for propulsion. In both cases however, the front legs have a brake effect on locomotion. To our knowledge, there are no study so far that measured the ground reaction forces (GRF) in loaded ants.

Nonetheless, it is possible to infer the propulsion behavior of the legs in our experiment based on the position of their tarsi. Indeed, as shown by Endlein and Federle (2015), depending on the GRFs, the tarsi attach differently to the substrate. The morphology of the tarsal attachment of *M. barbarus* (Figure S3) is comparable to that of other ants (Federle et al., 2001; Endlein and Federle, 2008). It seems thus fair to assume that they cling to the substrate in a similar way. As Endlein and Federle (2015), we observed in our videos two positions for the hind leg tarsi during the stance phase: on “heels”, during the first part of the stance phase, and on “toes”, during its second part (Figure S4). This would suggest that hind legs have a “compression and pushing” action in the first part of the stance phase, i.e. participate to propulsion, and then have a “tension and pulling” action on the last part of the stance phase, acting as a holding point for the ants not to fall over. For mid legs, the tarsi were usually in the “heel” position and were thus likely to participate in propulsion. These observations are purely qualitative as the resolution of our videos makes a quantitative analysis of these data tricky. The use of a miniature force plate (Bartsch et al., 2007; Reinhardt & Blickan, 2014b) to compare the GRFs of unloaded and loaded ants would provide crucial insights on how the different legs of the ants contribute to the stability and propulsion of loaded locomotion.

### Conclusion

We have shown in this study that unloaded *M. barbarus* workers display different gaits depending on their body mass. For big ants, these differences seem to be mainly explained by a compensation for the imbalance caused by their disproportionally bigger head. Small ants are able to carry proportionally heavier loads than big ants and scale effect provides a simple and satisfactory explanation for this difference. Moreover, our results show that loaded ants are often statically unstable during locomotion and that they maintain their balance by clinging to the ground. Further studies are required to determine the contribution of each leg to both stability and propulsion.

Big ants are more costly to produce than small ants. So why do colonies produce them if they are less efficient in transporting loads? One answer to this question is that, although big ants have lower load carriage performances than small ants, they are nonetheless able to carry on average loads of higher masses than small ants and to seize and transport items of larger diameters with their large and powerful mandibles (Fig. 3 in Bernadou et al., 2016). This could allow colonies to increase the size range of the food items retrieved to the nest so that they can enlarge their diet breadth and better match the size distribution of the food resources available in their environment (Davidson, 1978). Big ants may also play other roles than foraging in seed-harvesting ant colonies, such as removing the obstacles encountered on foraging trails, cutting thick plant stalks or milling the seeds inside the nest to prepare them for consumption. The significance of our results for the foraging ecology and division of labor in *M. barbarus* remains therefore to be investigated.

## Acknowledgements

The authors wish to thank Ewen Powie and Loreen Rupprecht who helped with data extraction, as well as Melanie Debelgarric who designed the Dufour gland extraction protocol.

## Author contributions

H.M., P.M. and V.F. conceived and designed the experiments and interpreted their results;

H.M. conducted the experiments and performed the analyses; V.F. and H.M. did the statistical analyses of the data; V.F. and H.M. wrote the paper and P.M. made essential contributions to the text; G.L helped for the collection of ant colonies and was in charge of their maintenance.

## Data availability

Dataset is available from http://doi.org/10.5281/zenodo.2646485

