## Supplementary information for "Walking kinematics in the polymorphic seed harvester ant *Messor barbarus:* influence of body size and load carriage"

#### I - Detailed Materials and methods

##### I – 0 – Vocabulary

For the sake of clarity, in the main text of our paper and in the supplementary material, we will refer to the (mesosoma + petiole) part of the ant as the "thorax" even if it contains some segments of the abdomen.

##### I – 1 – Computation of the ant CoM during displacement

The position of the center of mass (CoM) of the ants on each video frame was determined from the mass of their head, thorax and gaster and from the position of the CoM of these three body parts. To assess the positions of these CoMs, we tracked several points on each video frame (Figure 2) with the software Kinovea. We used the view of the top camera to determine the X and Y coordinates of the points (Figure 2 A-C) and that of the side camera (Figure 2 B-D) to determine the state of each leg during locomotion (i.e. in stance phase, swung or dragged).

On the top view, the following points were tracked on each video frame: the extremity of the mandibles, the neck, the junction between the petiole and the gaster and the extremity of the gaster. In addition, we tracked the extremity of the load carried by loaded ants (Figure 2). Assuming a homogeneous distribution of the mass within each body parts, we then computed for each frame the approximate (X, Y) position of the CoM of the three main body parts (plus the load) as the mean of the (X, Y) coordinates of the points located at their two extremities.

In addition, we used these tracked points in order to compute the length of the main body parts for each frame. The length of the main body parts was then computed as its average length over all frames of a video.

We weighed the ants at the end of each experiment to determine their total mass and separated each body parts to determine their mass. The ant CoM was then calculated as the barycenter of the CoM of the three main body parts weighed by their mass.

**I – 2 - Model for ideal tripod.**

Using R, we assessed the theoretical value of the static stability margin (SSM) for ants of different masses walking with an identical ideal alternating tripod gait.

In order to do so, we first modeled the horizontal displacement of the ant's CoM, depending on ant mass, and then the position of the supporting tripod. We used a coordinate system centered on the ant's neck, the X axis being the direction of walking motion and the Y axis the transverse direction (see Figure 2). We expressed all distances in units of body lengths.

We modeled the ant's CoM as the barycenter of the CoM of the ant's three main body parts. We considered that the CoM of these body parts in our local coordinate system have constant coordinates during locomotion. These coordinates depend on head length, thorax length and gaster length, which we were able to assess from the relationship between these lengths and ant mass in our data (Table S1). Finally, we calculated a theoretical mass for every body parts from the relationship between total body mass and the mass of each body parts in our data. This allowed us to compute the global CoM of the modelled ants.

We defined the ideal tripod gait as a perfect alternation of two tripods. Each leg thus has a duty factor of 0.5 and exactly three legs are in contact with the ground at any time. For the position of lift off and touch down of each leg, we took as a baseline the mean of the positions we observed in our experiments (Table 1). However, as leg length grows allometrically with ant mass (Table S2, Figure S6, Felden 2014), we corrected the position of lift off and touch down of ants of different masses based on the percentage deviance of its leg length from the mean leg length of ants calculated from a linear model (Table S2) (i.e. if its front leg was 20% longer than the average front leg in ants, the lift off and touch down positions were modified so that the X and Y coordinates of these positions were 20% higher). We then computed the theoretical position of the legs in the ant coordinate system at each time step as the linear interpolation between their touch down and lift off position, assuming a straight line for the trajectory of the legs between touch down and lift off.

Finally, we computed the SSM at each time step with the same method as that described in the Material & Method section of the paper, as well as its minimum value and the proportion of statically unstable locomotion over a walking cycle.

### II - Supplementary Tables

**Table S1: Relationship between total body mass (in mg) and each body part length and relative mass (N = 45).**

| Variable | Model prediction for mean(ant mass) ± CI | Slope for $log_{10}$(ant mass) ± CI | Adj $R^2$ |
|---|---|---|---|
| Head length (BL) | 0.277 ± 0.0047 | 0.0369 ± 0.0126 *** | 0.44 |
| Thorax length (BL) | 0.428 ± 0.0053 | -0.0376 ± 0.0141 *** | 0.39 |
| Gaster length (BL) | 0.297 ± 0.0056 | 0.0011 ± 0.0149 ns | -0.02 |
| Relative head mass | 0.385 ± 0.0071 | 0.1745 ± 0.0189 *** | 0.89 |
| Relative thorax mass | 0.254 ± 0.0067 | -0.0461 ± 0.0179 *** | 0.37 |
| Relative gaster mass | 0.293 ± 0.0108 | -0.0862 ± 0.0286 *** | 0.45 |

Each line gives the results of a linear model describing the relationship between $log_{10}$(ant mass) and the length of the three main body parts. The first column corresponds to the model prediction ± 95% confidence interval for the mean value of ant mass (i.e., 11.8 mg). The second column gives the coefficient of the model for $log_{10}$(ant mass) ± 95% confidence interval for (with the level of significance given by the t value: *** $p < 0.001$; ** $p < 0.01$; * $p < 0.05$; . $p < 0.1$; n.s. $p \geq 0.1$). The third column gives the adjusted $R^2$ for the linear model. BL= body length.

**Table S2: Relationship between body mass (in mg) and leg length (in mm)**

| Variable | Intercept ± CI | Slope for $log_{10}$(ant mass) ± CI |
|---|---|---|
| $log_{10}$(front leg length) (mm) | 0.276 ± 0.0073 *** | 0.310 ± 0.0075 *** |
| $log_{10}$(mid leg length) (mm) | 0.340 ± 0.0064 *** | 0.305 ± 0.0066 *** |
| $log_{10}$(hind leg length) (mm) | 0.459 ± 0.0075 *** | 0.283 ± 0.0075 *** |

Each line gives the results of a linear model describing the relationship between $log_{10}$(ant mass) and $log_{10}$(leg length). The first column corresponds to the model intercept ± 95% confidence interval. The second column gives the coefficient for $log_{10}$ (ant mass) ± 95% confidence interval (with the level of significance given by the t value: *** $p< 0.001$; ** $p < 0.01$; * $p < 0.05$; . $p < 0.1$; n.s. $p \geq 0.1$). The third column gives the adjusted $R^2$ for the linear model. Data from Felden (2014).

#### III - Supplementary Figures

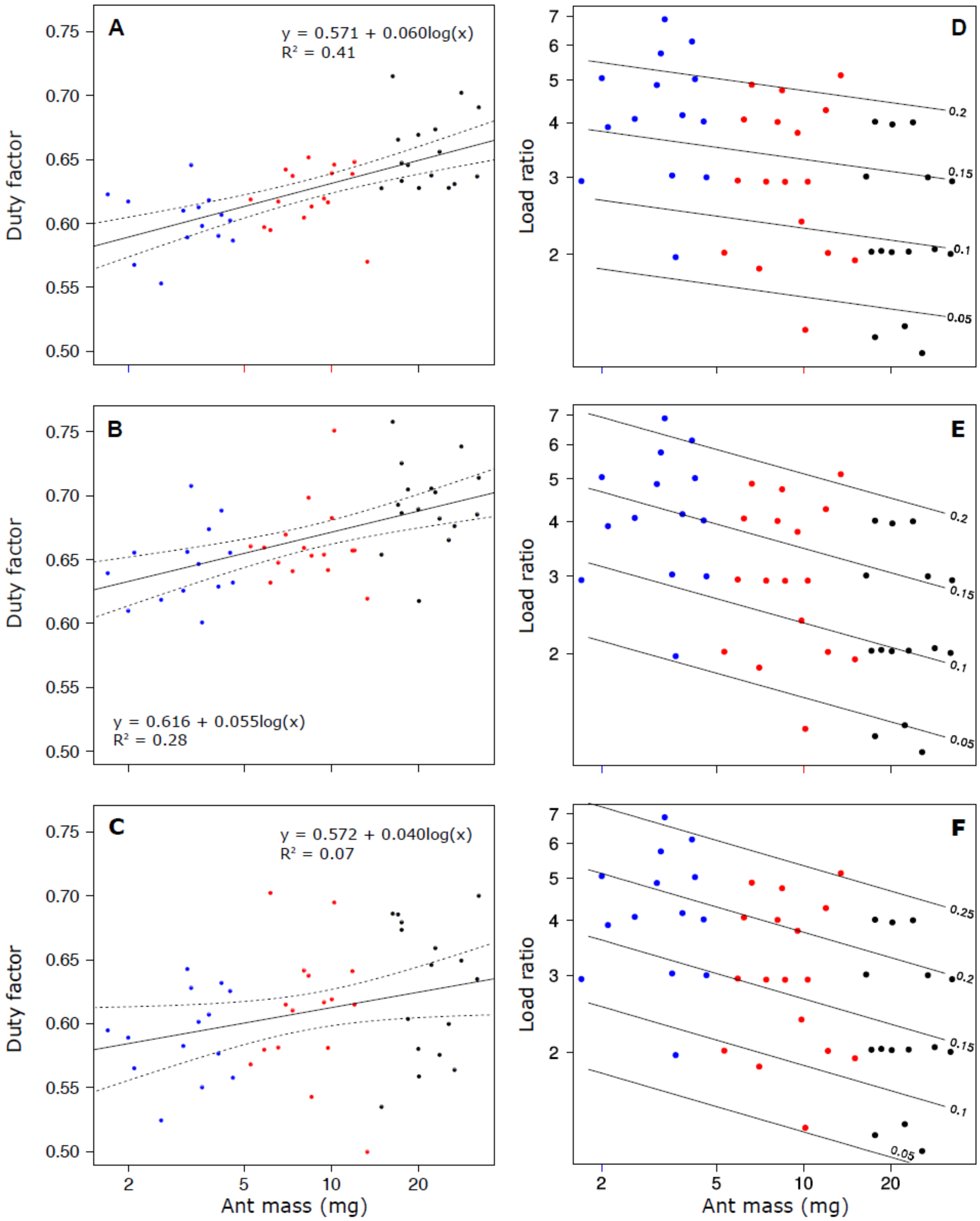


*Figure S1: **A-C: Duty factor as a function of ant mass** for front (**A**), mid (**B**) and hind (**C**) legs during unloaded locomotion. The straight lines give the predictions of a linear model and the dotted lines the 95% confidence interval of the slope of the regression line. **D-F: Change in duty factor as a function of ant mass and load ratio** for the front (**D**), mid (**E**) and hind (**F**) legs. The lines of equal changes in duty factor values are given by a general linear model. The points represent tested ants. N = 45 ants.*

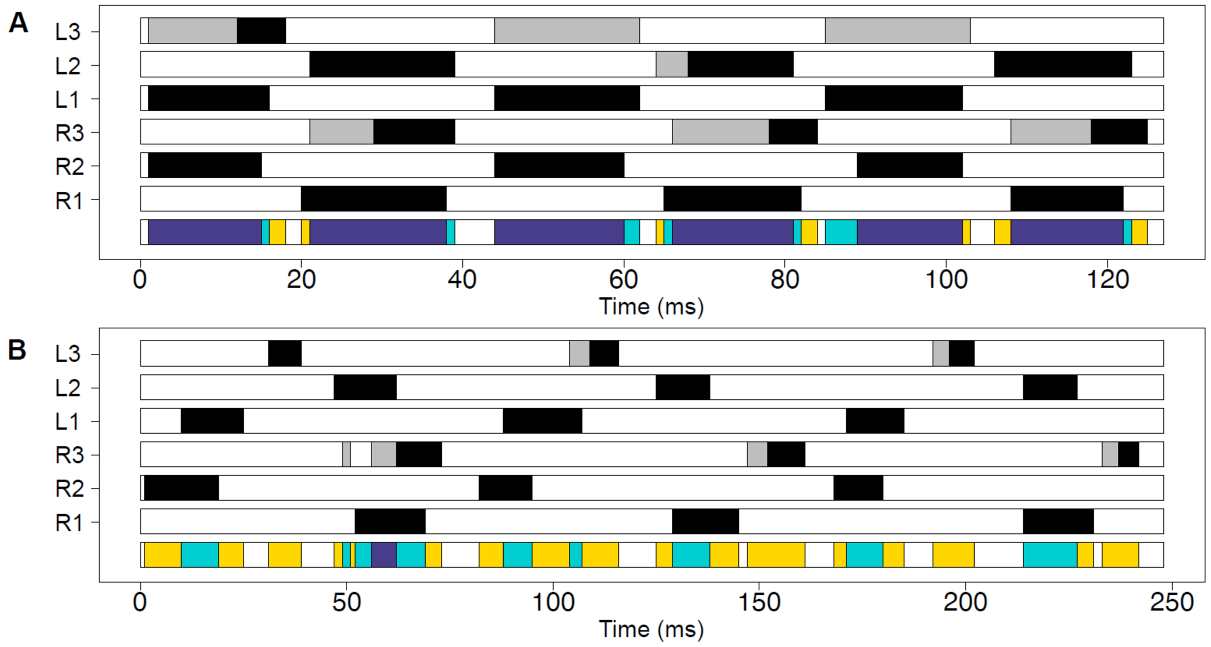


*Figure S2:* ***Example of inter-leg coordination*** *for one ant (ant mass = 3.1 mg) during unloaded (**A**) and loaded (**B**) locomotion (Load Ratio = 4.9). R1, R2, R3: right front leg, mid leg and hind leg; L1, L2, L3: left front leg, mid leg and hind leg. Black bars represent swing phases, white bars represent stance phases while grey bars represent dragging. The bottom line represents the number of legs in contact with the ground (including dragged legs): six (white), five (yellow), four (light blue) or three (purple).*

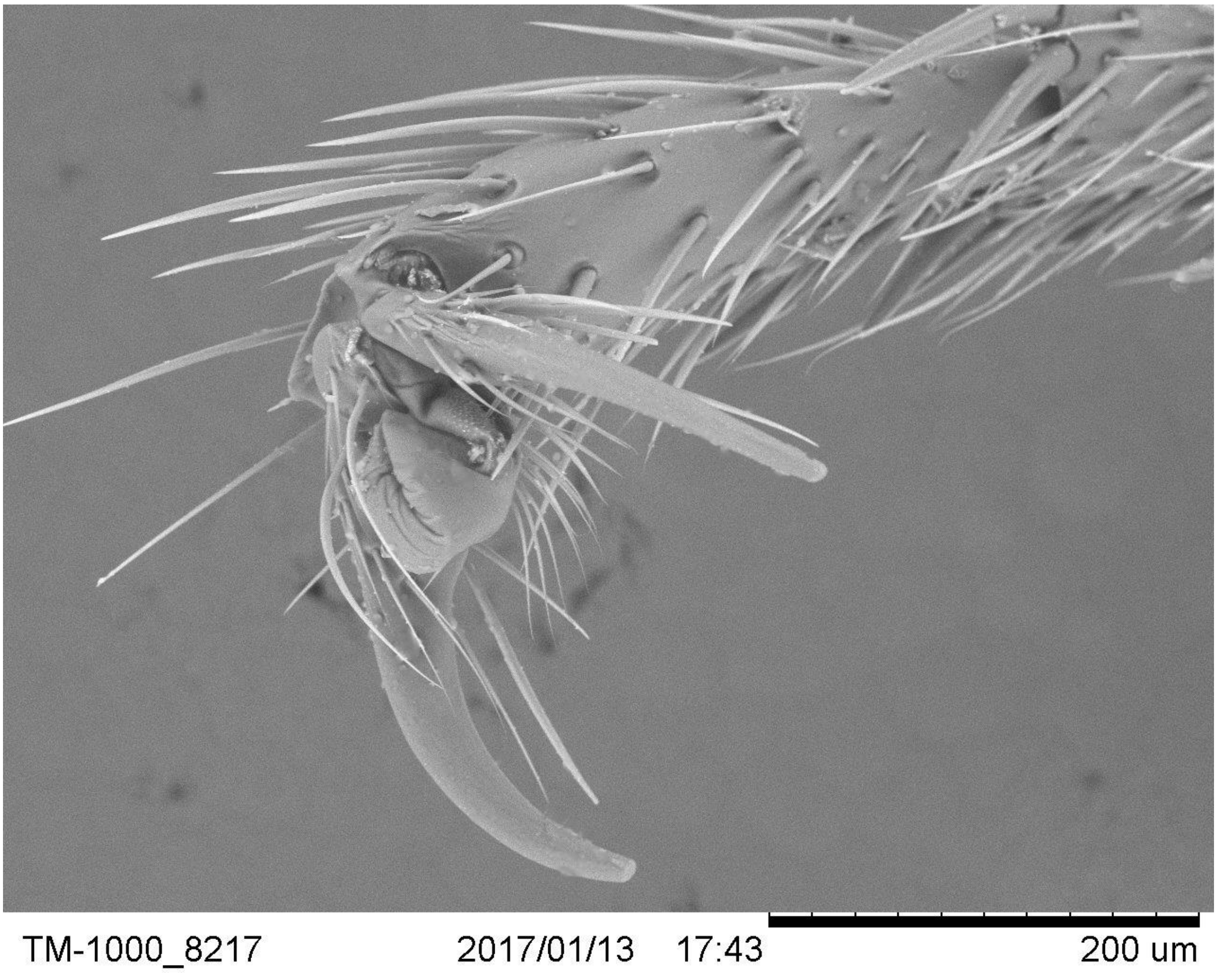


*Figure S3:* ***Scanning electron microscope*** *(Hitachi TM-1000 Tabletop Microscope)* ***image of a Messor barbarus hind leg tarsus showing the claws and the adhesive pad (arolium) between the claws****. Dry specimen was put without treatment in the microscope chamber.*

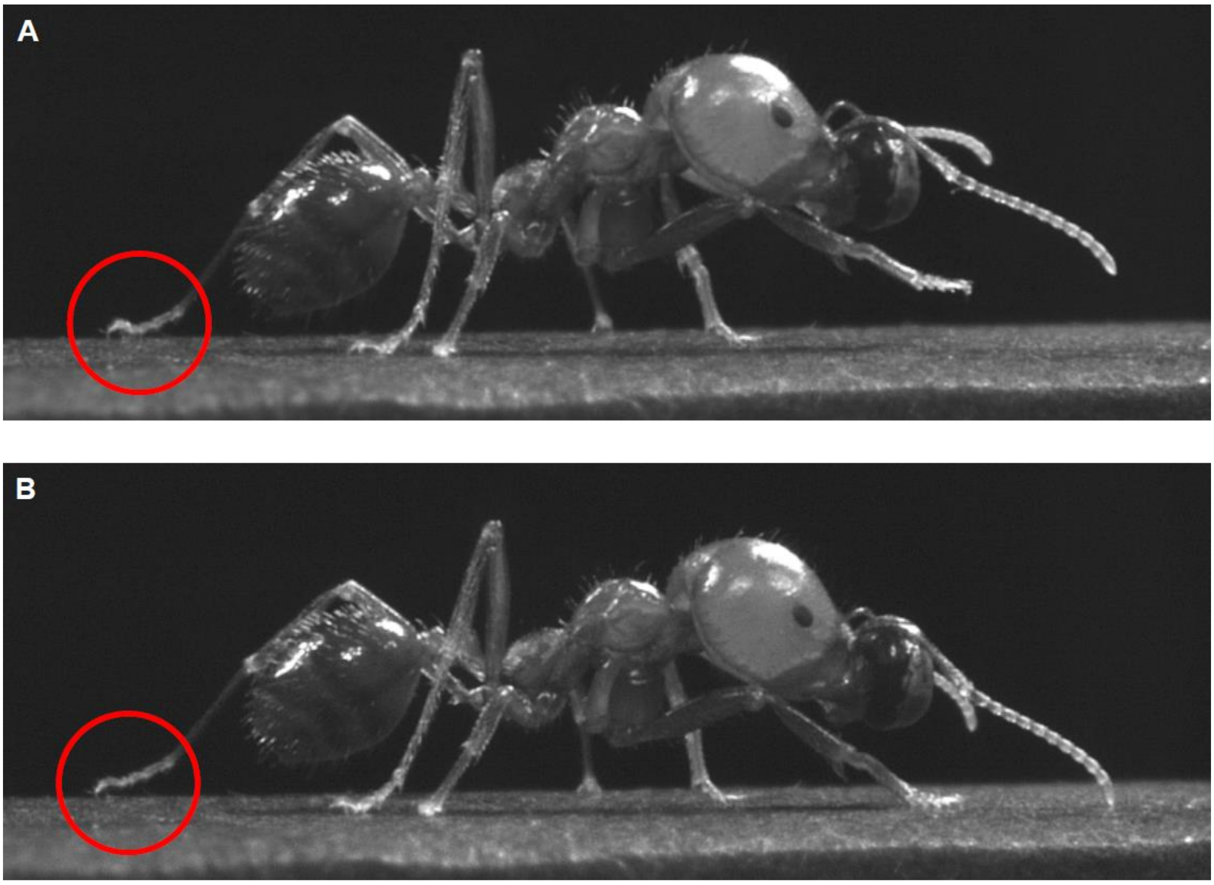


*Figure S4:* ***Left hind leg position during stance phase. A****: first part of the stance phase, the tarsi is on "heels";* ***B****: second part of the stance phase, the tarsi is on "toes".*

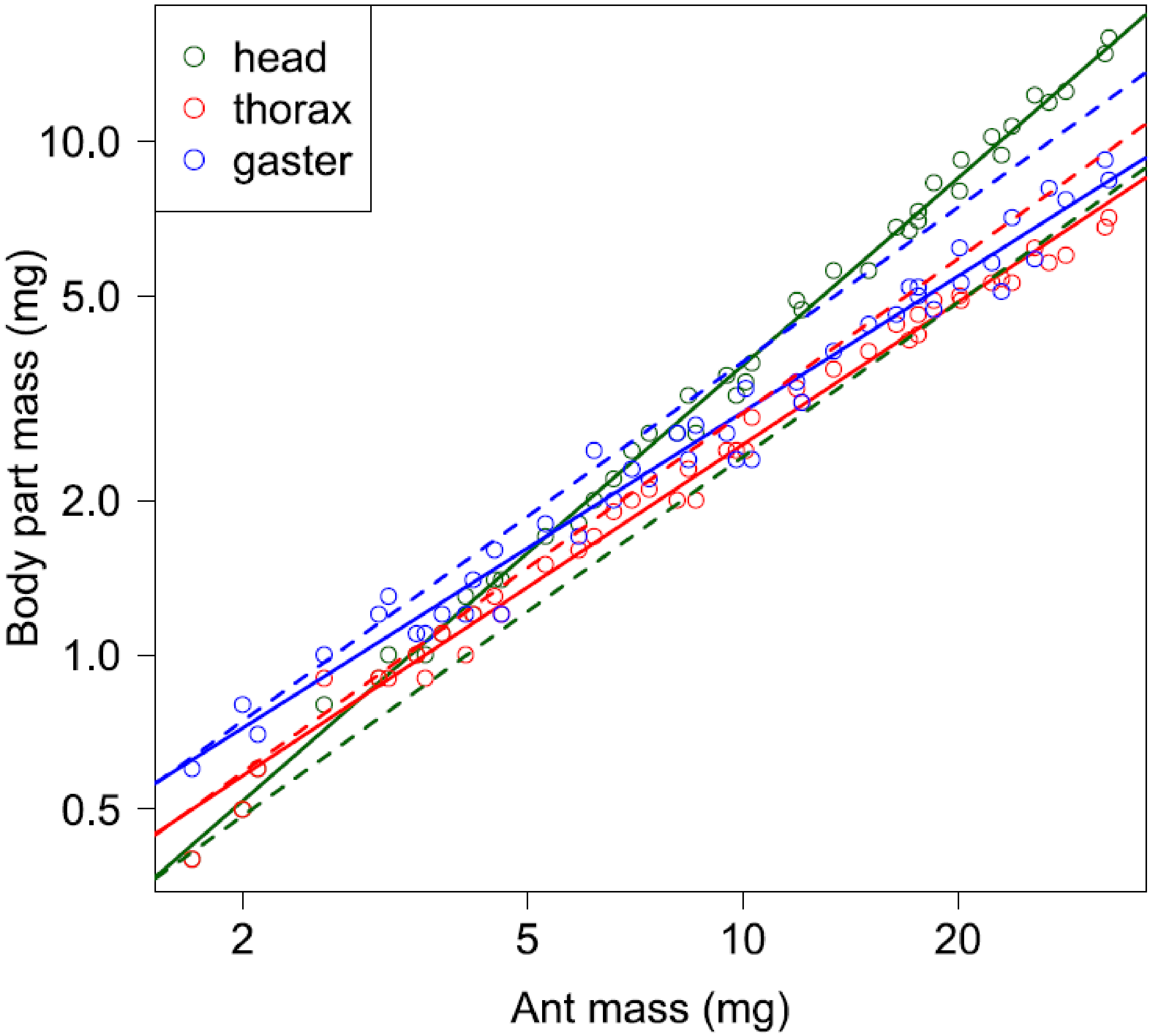


***Figure S5: Body part mass as a function of ant mass.*** *The solid lines represent regression model for head (green), thorax (red) and gaster (blue). Black dashed lines correspond to slope of 1 with unchanged elevation for each body part, i.e., what would be expected in absence of allometry.*

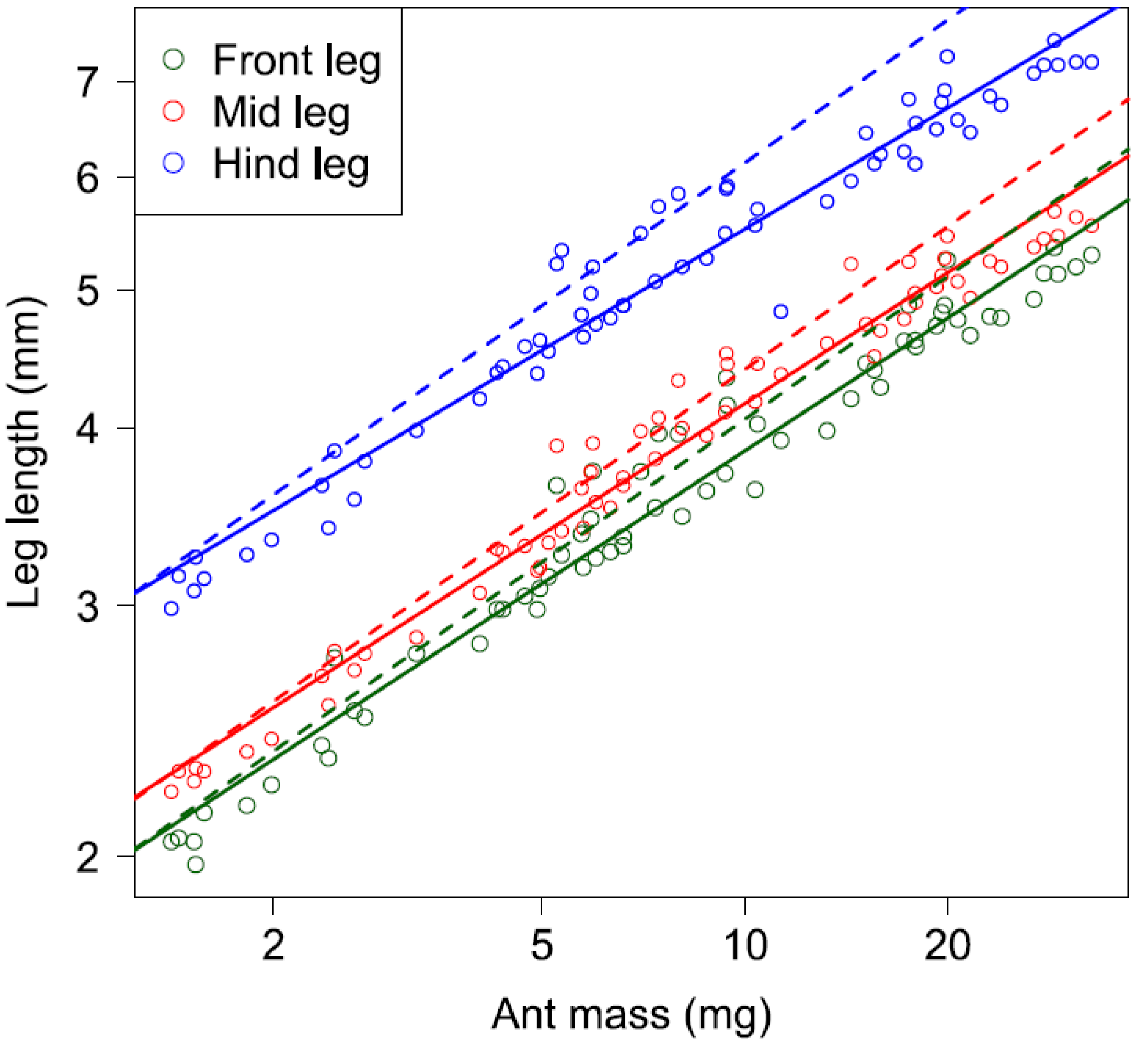


***Figure S6: Leg length as a function of ant mass.*** *The solid lines represent regression model for front (green), mid (red) and hind (blue) legs. The black dashed lines correspond to a slope of 1/3 with unchanged elevation for each leg, i.e. what would be expected if ants of different masses had geometrically similar shapes. Data from Felden (2014).*

#### IV - Movies

**Movie 1: Unladen ant walking.** Ant mass = 32.1 mg.

**Movie 2: Loaded ant carrying its load.** Ant mass = 32.1 mg. Load mass = 32.3 mg.

**Movie 3: Loaded ant pushing its load.** Ant mass = 15.1 mg. Load mass = 78.7 mg.
